# First Draft Genome of the Corn Leafhopper *Dalbulus maidis*: A Transparent, Open, and Updateable Resource in the Context of Agricultural Emergence

**DOI:** 10.1101/2024.06.25.600652

**Authors:** Humberto Julio Debat, Franco Fernández

## Abstract

The corn leafhopper, *Dalbulus maidis*, is a significant pest affecting maize crops, causing extensive economic losses and posing a threat to food security. This study presents the first draft genome of *D. maidis* as part of a comprehensive initiative to generate critical information for effective pest management and long-term control strategies. The genome sequencing is being conducted using a hybrid approach that integrates Oxford Nanopore Technologies (ONT) and Illumina platforms, ensuring high accuracy and depth. The initial genome assembly comprises approximately 580 Mb, with an N50 of 50,453 bp, indicating a draft assembly quality. The genome’s completeness, evaluated using BUSCO, stands at 68.6%, underscoring the thoroughness of the assembly. This first draft genome is designed to be a “living genome,” subject to continuous updates as new sequencing data become available. By providing an open and updatable genomic resource, this study aims to facilitate ongoing research and foster collaborative efforts in developing innovative solutions to mitigate the impact of *D. maidis* on maize cultivation.

## Introduction

*Dalbulus maidis*, commonly known as the corn leafhopper, holds profound agricultural significance throughout the Americas due to its role as a vector of maize pathogens (Virla., 2024). This insect species is notorious for transmitting pathogens maize bushy stunt phytoplasma and corn stunt spiroplasma associated with corn stunt disease complex and viruses, which can devastate maize crops (Nault., 1998). Its ability to spread these pathogens efficiently across vast agricultural regions poses a significant threat to maize production, a staple food crop crucial for food security and economic stability across the continent (Oliveira et al., 2023).

Corn plantations across the southern regions of Brazil, the northern regions of Argentina, and recently within the core zones of the latter have been increasingly threatened by the corn leafhopper. This pest has significantly proliferated, driven in part by climate change, which has created favorable conditions for its spread and survival (Santana et al., 2019). The economic implications of this pest infestation are severe, with estimates indicating losses exceeding US$2.2 billion in the last season alone in Argentina (Vazquez., 2024), due to crop damage and the transmission of associated diseases such as phytoplasma, corn stunt spiroplasma and maize rayado fino virus (Nault et al., 1980).

The importance of *D. maidis* as a pest lies not only in its direct impact on maize yield but also in its role as a vector for several phytopathogenic viruses and phytoplasmas (Oliveira et al., 2023). These pathogens exacerbate the damage caused by the leafhopper, leading to complex disease dynamics that are challenging to manage (Atmaca et al., 2014). Current control measures, primarily based on chemical insecticides and cultural practices, have proven insufficient and unsustainable in the long term (Tsai et al., 1990; Neves et al., 2022).

Given this critical scenario, it is imperative to generate strategic information that supports both immediate and long-term pest management solutions. Genomic information provides a powerful tool for understanding pest biology, ecology, and evolution, enabling the development of innovative and sustainable management strategies (Poelchau et al., 2016). However, there is a notable lack of genomic resources for *D. maidis*, which hampers efforts to devise effective control strategies (Frey et al., 2022).

In response to this need, the National Institute of Agricultural Technology (INTA) in Argentina has initiated a project to sequence, assemble, and annotate the genome of *D. maidis*. This work presents the initial draft genome assembly of *D. maidis*, providing the first set of massive sequences and a detailed chronology of the sequencing process. This dynamic draft genome is expected to serve as a foundational resource for future research and development in pest management strategies, contributing to the sustainability of maize cultivation.

## Materials and Methods

### Insect Colony

Specimens of *Dalbulus maidis* used in this analysis were sourced from a healthy colony maintained at the Institute of Plant Pathology (CIAP-INTA) in Córdoba, Argentina in May 2024. The colony was propagated under controlled greenhouse conditions, with a consistent temperature of 25°C and a relative humidity of 60%. The management of the colony were overseen by Mariana Ferrer and Karina Torrico, ensuring the health and viability of the specimens.

### Nucleic Acid Extraction

For the extraction of high molecular weight genomic DNA (gDNA), the Monarch® HMW DNA Extraction Kit for Tissue (NEB, USA) was utilized in accordance with the manufacturer’s specifications. Twenty male specimens of *Dalbulus maidis* were homogenized in liquid nitrogen using a mortar and pestle to ensure complete tissue disruption. The quality and quantity of the extracted gDNA were assessed using several methods to ensure suitability for genomic analyses. Spectrophotometry (Nanodrop-1000, Thermo Scientific) was employed to measure the purity and concentration of the DNA, providing a 260/280 ratio to indicate protein contamination. Fluorometry (Quantus, Promega) was used for accurate quantification of the gDNA, which is essential for downstream applications. Additionally, the integrity of the gDNA was verified through electrophoresis in 1% agarose gels stained with GelRed® Nucleic Acid Gel Stain (Biotium, USA), allowing visualization of high molecular weight DNA and detection of any degradation. This comprehensive evaluation ensured that the gDNA met the high standards required for subsequent sequencing processes, contributing to the robustness and accuracy of the genome assembly.

### Sequencing using a Hybrid ONT + Illumina Strategy

For the initial sequencing stage, 1.2 µg of high molecular weight genomic DNA (gDNA) was utilized to construct libraries on the Oxford Nanopore Technologies (ONT) platform, specifically for long-read sequencing. The Ligation Sequencing Kit (SQK-LSK109) was employed in accordance with the manufacturer’s instructions. The resulting library was quantified using fluorometry (Quantus, Promega) to ensure accurate measurement of the DNA concentration. Sequencing was then performed on a MinION device with a 1Kb flowcell and R9.4.1 chemistry at the IPAVE-CIAP facility in Córdoba, Argentina. The raw sequencing data were processed using the Dorado software (version 0.7.1) in sup mode for basecalling, which converts raw signal data into nucleotide sequences. This initial stage focused on generating long reads that are essential for constructing a robust genome assembly. In the second sequencing stage, 1 µg of the same gDNA is being sequenced using the Illumina NovaSeq platform. This stage will employ the 150PE TruSeq DNA Nano 350bp library preparation kit to generate paired-end reads, which will complement the long reads from the ONT platform and provide a comprehensive genomic dataset. This hybrid sequencing approach leverages the strengths of both technologies, combining the long-read capabilities of ONT with the high accuracy and depth of coverage provided by Illumina sequencing.

### De novo Assembly

The reads obtained from the first stage of sequencing using the ONT platform were processed to ensure high-quality data for downstream analysis. Adapter sequences were trimmed using Porechop version 0.2.4, and the reads were filtered based on quality (--min_mean_q 15) and length (--min_length 1000) using Filtlong v0.2.1. These preprocessing steps are crucial for removing low-quality data and ensuring that only high-quality reads are used for assembly. The primary statistics of the reads, both pre- and post-trimming, were evaluated using NanoComp version 1.19.1, providing a comprehensive overview of read quality and distribution. Following preprocessing, the filtered reads were assembled de novo using Flye v2.9.1 on the local CIAP-INTA HPC server. The assembly parameters included -- nano-hq fastq, --genome-size 1g, -t 20, --read-error 0.03, and --meta -i 4. The genome size was estimated based on previous reports of leafhoppers, while the -- meta option allowed for the assembly of mitochondrial sequences and potential symbiont organisms such as bacteria and viruses. Post-assembly, the genome was polished using Medaka version 1.12.0 with the r941_min_sup_g507 model to correct base-call errors and enhance the overall assembly quality. General assembly metrics, including contig length and N50 values, were evaluated using Quast version 5.2.0 on the usegalaxy.eu platform, providing insights into the completeness and accuracy of the assembly. Additionally, the completeness of the genome assembly was assessed using BUSCO (Benchmarking Universal Single-Copy Orthologs) with the insecta_odb10 lineage dataset, as described by Seppey et al. (2019). This assessment ensures that the genome assembly contains a comprehensive representation of conserved single-copy orthologs, which is indicative of the assembly’s completeness and utility for further biological studies.

### Data availability

The raw ONT long raw reads have been deposited at the NCBI Sequence Read Archive (SRA) under the Bioproject PRJNA1120083 (BioSample accession SAMN41674197). The *Dalbulus maidis* genome assembly has been shared publicly through the INTA institutional digital repository under accession http://hdl.handle.net/20.500.12123/18043, guaranteeing that this resource is available to the entire scientific community. The assembly will be updated continuously whenever additional data is available, which will be processed in real time.

## Results and Discussion

### Quality of High Molecular Weight gDNA

The extracted genomic DNA (gDNA) met the stringent quality requirements necessary for high-throughput sequencing applications. The purity of the DNA was confirmed by a 260/280 ratio of 1.97, as measured by a Nanodrop spectrophotometer, indicating minimal contamination with proteins or other impurities. Additionally, the integrity of the gDNA was verified by agarose gel electrophoresis, where an intense, high molecular weight DNA band was observed without signs of degradation. The final concentration of the gDNA was accurately quantified using the Quantus fluorometer, yielding a concentration of 116 ng/µl.

### Sequencing and Assembly

Over a continuous 48-hour period, the sequencing run included a library reload at approximately 24 hours to optimize flow cell performance. Initially, 2,397,865 raw reads (totaling 6,690,474,495 bases) were generated, featuring an N50 of 5,823 bp and an average quality score of 14.1. Post-processing, 1,398,710 reads (6,151,663,624 bases total) were retained, showing improvements with an N50 of 6,782 bp and an average quality score of 15.1. The resulting assembly generated with Flye (Dalbulus_CIAPv0.1 version) comprised 22,107 contigs, encompassing a final size of 584,127,104 bases (approximately 580 Mb), with an N50 of 50,453 bp (refer to Table 1). Completeness was assessed using BUSCO (mode genome) against the insecta_odb10 lineage (comprising 75 genomes and 1,367 BUSCOs), the analysis yielded a 68.6% completeness score. The Dalbulus_CIAPv0.1 genome assembly serves as a valuable initial resource for further research into insect genomics. However, it should be noted that this assembly represents a preliminary draft, and significant improvements can be achieved through additional sequencing efforts. Future iterations should incorporate new flow cell runs using both long-read sequencing technologies like Oxford Nanopore and short-read technologies such as Illumina. These advancements, already in course, are expected to enhance the assembly’s contiguity, completeness, and overall quality metrics, particularly in terms of BUSCO assessment. By refining these genomic metrics, the assembly will more accurately reflect the complexities and nuances of the actual insect genome, facilitating deeper insights into evolutionary processes, genetic adaptations, and ecological interactions.

**Table 1:** General statistics of the *Dalbulus maidis* v0.1 INTA partial genome assembly.

| Parameter | Dalbulus_CIAPv0.1 |
| --- | --- |
| Contigs | 22.107 |
| Total length | 584.127.104 |
| Largest contig | 813.079 |
| N50 | 50.543 |
| N90 | 11.594 |
| GC% | 34.78% |
| Ns per 100kbp | 0 |
| BUSCO | 68.6% |

### Perspectives

The emergency facing the agricultural sector demands not only immediate management actions but also a forward-looking strategic approach that fosters innovative solutions. The genome of the *Dalbulus maidis* insect represents an invaluable and essential resource for understanding and addressing the pathosystem affecting the maize chain, which has significant implications for the agricultural sector. It is crucial to emphasize that this is an ongoing process of refinement. The initial draft of the *Dalbulus maidis* genome (version 0.1) generated in this work will evolve continuously, designated as a “living genome,” with regular updates as new sequencing data become available. These updates will incorporate advancements from ongoing sequencing efforts, including both emerging technologies like Oxford Nanopore and established methods such as Illumina short reads. Raw sequencing data is being promptly deposited in the NCBI Sequence Read Archive (SRA) under the Bioproject PRJNA1120083 (BioSample accession SAMN41674197). This commitment by INTA underscores our dedication to transparency and accessibility in scientific data, facilitating secondary analyses and fostering community-driven knowledge generation through immediate data release.

---

**Figure 1.**
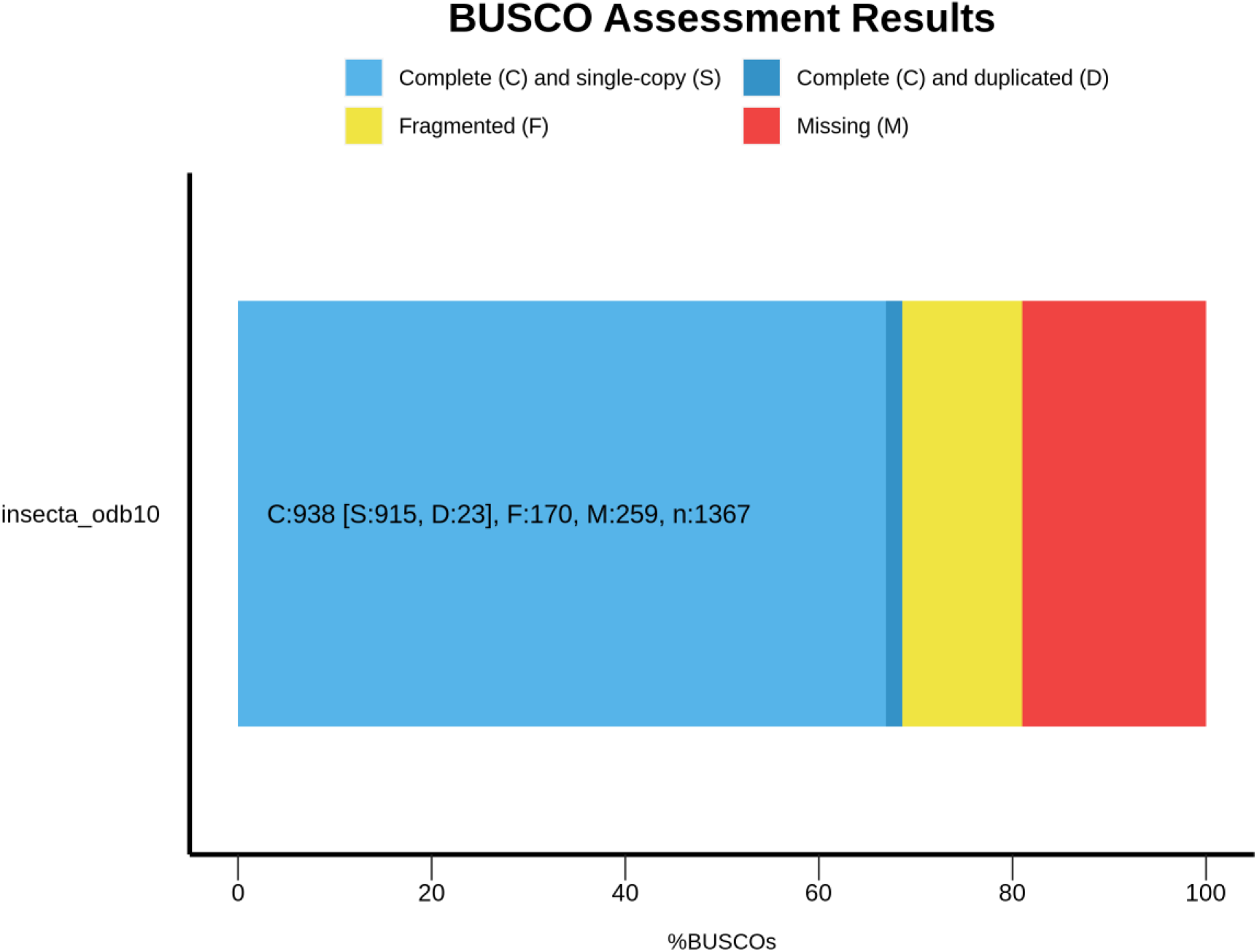
BUSCO (mode genome) analysis values obtained with the *Dalbulus maidis* v0.1 INTA partial genome using the insecta_odb10 lineage (number of genomes: 75, number of BUSCOs: 1367)

